# Gaze shifts in freely moving mice comprise distinct head-eye coordination motifs

**DOI:** 10.64898/2026.06.13.732086

**Authors:** Yuchen Hou, Marius Schneider, Jhoseph Shin, Cristopher M. Niell, Michael Beyeler

## Abstract

Freely moving mice are generally thought to redirect gaze mainly through head-coupled movements, with eye movements stabilizing the retinal image or resetting eye position. This reflexive view leaves unresolved whether gaze shifts also include active coordination modes. Here we show that a single gaze-shift class resolves into multiple distinct head-eye coordination motifs. Using high-resolution head and eye tracking in freely moving mice, we identified four reproducible motifs: Head-before-Eye (HbE), Head-with-Eye (HwE), Head-Dominant (HD), and Eye-Dominant (ED). The motifs differed in head-eye timing, locomotor context, and pre-onset visual-behavioral structure. Their short-timescale sequential organization was non-random: HD formed alternating side-to-side head movements, whereas HwE-HbE-ED formed a directed transition chain preserved within each movement direction. Simultaneous neural recordings further revealed motif-dependent responses in primary visual cortex (V1) and superficial superior colliculus (sSC), with stronger sSC than V1 modulation for the three motifs distinguished by pre-onset visual-behavioral structure (all but HbE). HwE showed the clearest signature of active orienting, combining near-synchronous head-eye onset, strong relevance to visual-behavioral features, and the largest pre-onset and earliest peri-onset responses in sSC. These results expand a unitary view of gaze shifts in freely moving mice into a diverse, structured repertoire of coordination motifs, and identify HwE as a candidate active gaze-shift mode during natural behavior.

## 1 Main

Gaze control is a fundamental component of active vision. Animals move their eyes, head, and body to sample the environment, alternating rapid gaze shifts with intervening periods of stabilization in a “saccade and fixate” strategy (Land and Tatler, 2009 ; Yarbus, 1967 ; Land, 1999).

In foveated species, gaze shifts are coordinated head-eye movements that redirect the fovea toward informative regions of a scene (Land and Tatler, 2009 ; Krauzlis et al., 2013 ; Hayhoe and Ballard, 2005). In mice and other afoveated animals, eye movements are more tightly coupled to head and body motion (Wallace et al., 2013 ; Land, 2019 ; Meyer et al., 2020 ; Ambrad Giovannetti and Rancz, 2024 ; Skyberg and Niell, 2024 ; Parker et al., 2022a). These eye movements have been traditionally interpreted as stabilizing the retinal image during head motion rather than actively redirecting gaze toward targets. In this view, the vestibulo-ocular reflex (VOR) and optokinetic reflex (OKR) rotate the eyes counter to head motion to hold the scene stable on the retina, followed by brief eye movements to reset eye position in the orbit (Meyer et al., 2020).

Freely moving mouse gaze is usually analyzed within this framework, using the head-unrestrained head–eye terminology (McCluskey and Cullen, 2007 ; Verdone et al., 2026). Gaze speed is defined as the sum of eye and head rotational speed (Parker et al., 2023), thereby representing the net movement of the eyes relative to the scene. Gaze shifts are detected as brief, high-speed head–eye events in which the eyes move with the head, distinct from the slower stages in which the eyes counter-rotate against head motion (Meyer et al., 2020). Similar classification has been used to study prey capture (Michaiel et al., 2020), visual pursuit (Holmgren et al., 2021), free exploration (Parker et al., 2023; Sharp et al., 2025), and gaze-related responses in primary visual cortex (V1) and superior colliculus (SC) (Parker et al., 2023; Sharp et al., 2025). However, this approach treats gaze shifts as a single event class, usually defined by instantaneous head–eye velocity. This does not resolve whether the gaze shift class itself contains substructure and whether natural mouse gaze shifts are one reactive motor pattern or a repertoire of coordination modes with different sensory and neural signatures.

A growing body of work suggests that the unitary view is incomplete, and that mice possess the circuitry for active gaze control. Stimuli can evoke saccades coincident with attempted head rotations (Zahler et al., 2021). Gaze shifts depend on SC activity (Zahler et al., 2021, 2023; Wang et al., 2015; Gandhi and Katnani, 2011), modulate V1 responses (Parker et al., 2023 ; Miura and Scanziani, 2022), and can support visually guided task behavior (Itokazu et al., 2018 ; Sato et al., 2019 ; Sharp et al., 2025 ; Holmgren et al., 2021). Body-fixed experiments have also revealed head–eye co-initiation patterns that resemble active orienting (Verdone et al., 2026). This evidence indicates that mouse gaze is not purely reflexive stabilization, and that a strategy for active head–eye coordination may operate in mice (Verdone et al., 2026). What remains unresolved is whether such coordination modes appear during natural freely moving behavior, where gaze shifts have largely been treated as a single head-coupled category (Ambrad Giovannetti and Rancz, 2024).

A single gaze-shift label is also at odds with what has emerged in other visual systems. In larval zebrafish, saccades that appeared to form one class separated into multiple kinematic types used in different behavioral contexts (Dowell et al., 2024). A similar structure has been reported in other afoveated species (Salem et al., 2020 ; Mongeau and Frye, 2017). The same may be true in mice: head-coupled gaze shifts could contain multiple coordination motifs with different relationships to behavioral and neural activity.

Here, we combined high-resolution eye and head tracking (Fig. 1A) with simultaneous recordings from V1 and SC in freely moving mice. We asked whether natural gaze shifts form a single behavioral mode or a set of time-resolved head– eye coordination motifs. Unsupervised partitioning of perievent head–eye kinematics revealed four reproducible motifs (Fig. 1E): Head-before-Eye, Head-with-Eye, Head-Dominant, and Eye-Dominant. These motifs differed in head–eye timing, locomotor context, short-timescale sequence structure, preonset visual-behavioral structure, and peri-event modulation in V1 and SC. Head-with-Eye was the clearest candidate for active orienting: it combined near-synchronous head–eye onset, visual-behavioral predictability, and early sSC modulation. Together, these results show that natural mouse gaze control is not a single reactive pattern but contains structured coordination motifs linked to distinct behavioral and neural dynamics.

**Fig. 1.**
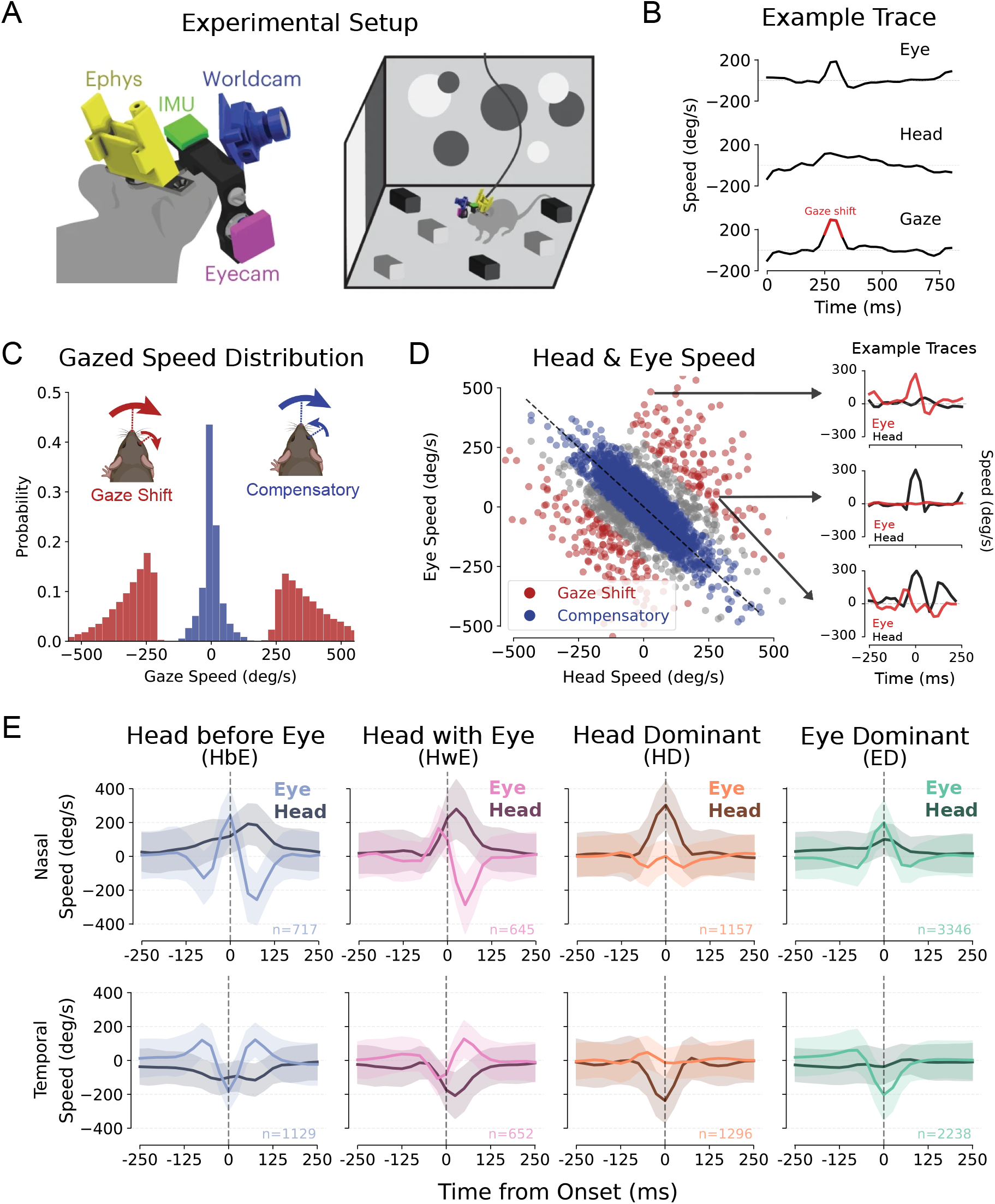
Time-resolved classification of head–eye movements in freely moving mice. **A,** Head-mounted recording system combining electrophysiology, an inertial measurement unit (IMU), a forward-facing world camera, and an infrared eye camera during open-field exploration (reproduced from Parker et al. (2023)). **B,** Example eye, head, and resulting gaze velocity traces during a gaze shift. Red indicates the detected gaze-shift event. **C,** Distribution of gaze velocities for gaze shifts and compensatory movements (insets reproduced from Parker et al. (2023)). **D,** Instantaneous eye velocity plotted against head velocity. Compensatory movements occupy the diagonal head–eye velocity axis, whereas gaze shifts span a broader region of the velocity space. Insets show example peri-event head and eye velocity traces. **E,** Mean peri-onset head and eye velocity traces for four gaze-shift motifs identified from time-resolved head–eye kinematics, shown separately for nasal and temporal movements. Shading indicates standard deviation across events; event counts are shown in each panel.

## 2 Results

### 2.1 Time-resolved head-eye kinematics reveal four gaze-shift motifs

In freely moving mice, eye movements have generally been divided into two broad classes: compensatory movements, in which eye motion opposes head rotation to stabilize visual input, and gaze shifts, in which head and eye motion combine to redirect gaze (Meyer et al., 2020 ; Parker et al., 2023). Consistent with prior work using this dataset and framework, instantaneous head–eye velocity separated compensatory movements from gaze shifts (Fig. 1A–D). However, this instantaneous classification treats all gaze shifts as one event class.

This operational gaze-shift class contained substantial temporal heterogeneity. Individual events varied in whether the head or eye contributed more strongly to the movement, and events with similar instantaneous gaze velocity could have markedly different peri-event trajectories (example traces in Fig. 1D). We therefore asked whether natural gaze shifts contain reproducible head–eye coordination motifs that are not visible from a single time point.

We represented each gaze-shift event by its peri-event kinematic profile over a *±* 100 ms window around movement onset, using horizontal head velocity, eye velocity, head acceleration, and eye acceleration sampled across the window. After *z*-scoring these features, we clustered events separately for each movement direction, then matched the resulting motifs across nasal and temporal gaze shifts (Fig. 1E; Methods 4.3). Because freely moving gaze shifts occupy a continuous movement space, we treated clustering as a compact decomposition into dominant coordination motifs, not as evidence for sharply separated behavioral categories.

We selected a four-motif solution because it balanced interpretability and reproducibility. At lower values of *k*, head-before-eye and near-synchronous head–eye movements were merged despite being separable in restrained preparations (Zahler et al., 2021; Verdone et al., 2026). At higher values of *k*, additional clusters mainly subdivided existing temporal profiles without producing qualitatively new head–eye dynamics (Fig. S1). The four-motif solution remained stable across animals in leave-one-mouse-out validation (Fig. S2) and recovered matched motifs in nasal and temporal gaze shifts, with cross-direction classification well above chance (Fig. S3). Thus, the four motifs provide a reproducible, literature-aligned summary of the dominant temporal structure within the gaze-shift class.

We refer to these motifs by their dominant coordination pattern: Head-before-Eye (HbE), Head-with-Eye (HwE), Head-Dominant (HD), and Eye-Dominant (ED) (Fig. 1E). The HbE motif was characterized by an initial head movement followed by eye movement in the same direction (16.51% of events). The HwE motif showed near-synchronous head and eye movement at onset, followed by a compensatory movement in the opposite direction (11.60% of events). The HD motif was driven primarily by head motion, with comparatively limited eye movement (21.94% of events). The ED motif showed the converse pattern: a large eye movement with limited accompanying head motion (49.94% of events) ^1^.

The operational gaze-shift class recovered by instantaneous head–eye velocity analysis comprised multiple time-resolved coordination motifs, providing the basis for subsequent analyses of locomotor context, visual-behavioral input, and neural activity.

### 2.2 Gaze-shift motifs occur in distinct locomotor and sequential contexts

All four motifs were interleaved during free exploration rather than restricted to separate recording epochs (Fig. 2A). Given that the four motifs have similar gaze speed (Fig. S4), we first confirmed that the pooled motifs retained the head–eye coordination structure evident in the partitioning. For each event, we computed the head-minus-eye speed difference in two pre-onset windows, from *−*100 ms to *−*50 ms and from *−*50 ms to onset (Fig. 2B). Positive values indicate greater head speed; negative values indicate greater eye speed. The motifs separated along the expected coordination axes: HbE events were head-biased before onset, HwE events showed little pre-onset imbalance, HD events became strongly head-dominated immediately before onset, and ED events showed the opposite pattern, with eye speed exceeding head speed near onset.

**Fig. 2.**
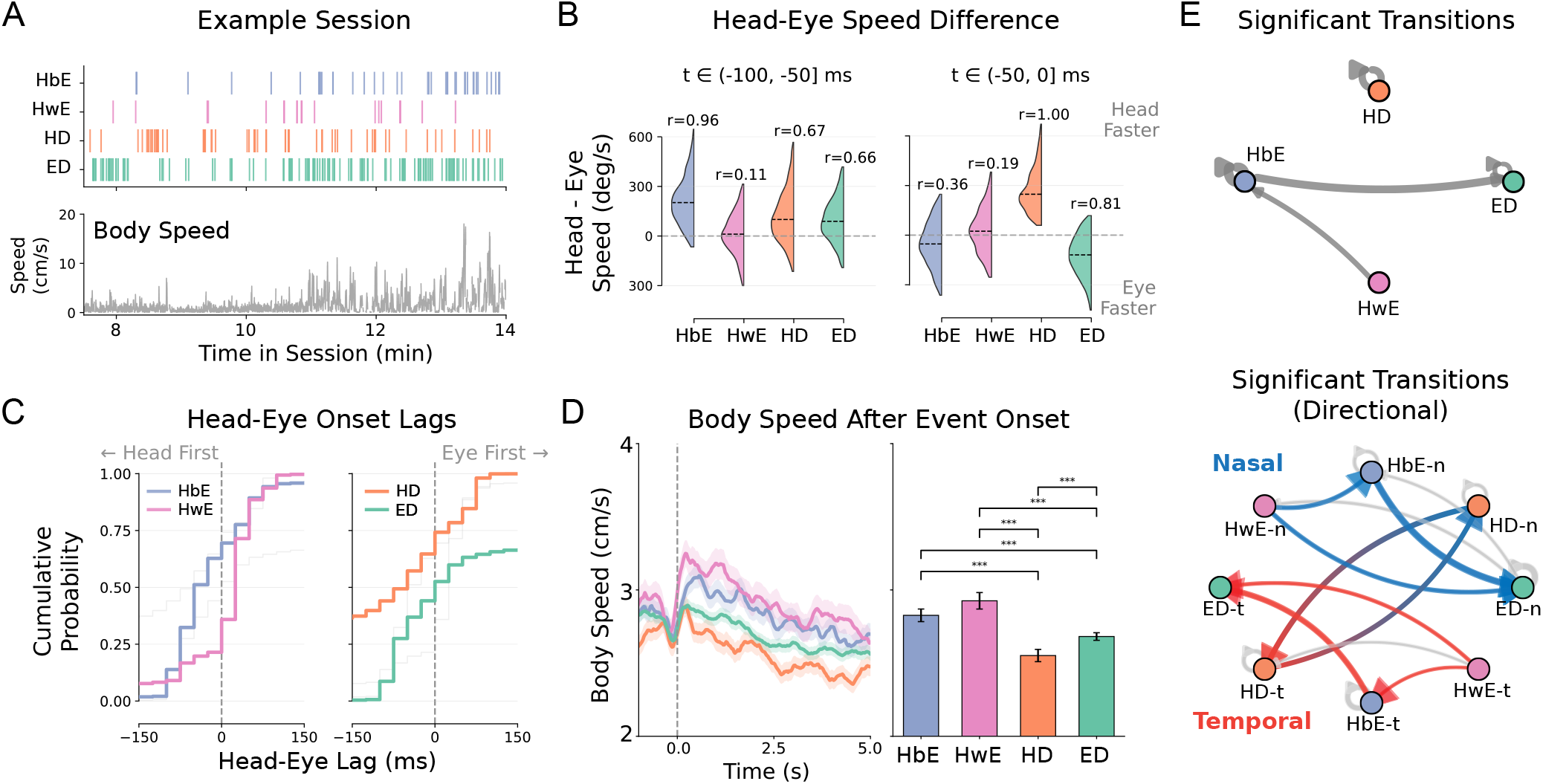
Gaze-shift motifs occur in distinct locomotor states and structured sequences. **A,** Example freely moving session showing gaze-shift events colored by motif, aligned with the animal’s body speed. **B,** Head-minus-eye speed difference for each motif in two pre-onset windows: *−*100 ms to *−*50 ms and *−*50 ms to onset. Positive values indicate greater head speed; negative values indicate greater eye speed. Half-violins show event distributions. Rank-biserial correlations (*r*) report effect sizes for one-sample Wilcoxon signed-rank tests against zero. **C,** Cumulative distributions of head–eye onset lag for each motif. Lag was defined as head onset minus eye onset, so negative values indicate head-first events and positive values indicate eye-first events. **D,** Body speed aligned to gaze-shift onset for each motif. Left, mean *±* s.e.m. body-speed trajectories from *−*1 s to +5 s relative to onset. Right, mean *±* s.e.m. body speed averaged over the 5 s post-onset window. Asterisks indicate significant pairwise comparisons after FDR correction. **E (top),** Pooled short-timescale transition structure among gaze-shift motifs. Edges indicate significantly enriched transitions within 500 ms of each gaze shift, relative to the expected probability given the marginal frequency of the target motif (*z >* 1.96, permutation test). HD forms a recurrent self-transition, whereas HwE, HbE, and ED form an asymmetric transition chain. **E (bottom),** Direction-resolved transition graph for nasal (“-n”, blue) and temporal (“-t”, red) gaze shifts. Nodes indicate motif–direction states; edges indicate significantly enriched transitions within 500 ms. HD provides the dominant transitions between nasal and temporal directions, whereas the HwE–HbE–ED chain is preserved within each movement direction.

Head–eye onset timing showed the same organization. For each event, head and eye onset were defined as the first time point at which each velocity trace exceeded 50% of its perievent peak. Head–eye lag was defined as head onset time minus eye onset time, so negative values indicate head-first events and positive values indicate eye-first events (Fig. 2C; Methods 4.4). The onset-lag distributions mirrored the speed-difference analysis. HbE and HD events were predominantly head-first, HwE events were mostly simultaneous or eye-first, and ED events were mostly eye-first. Thus, the motifs remained separable by both the relative speed and timing of head and eye contributions.

The motifs were also associated with different locomotor dynamics (Fig. 2D). HwE yielded the highest post-shift locomotion speed, while HD had the lowest locomotion speed, and HbE and ED were in-between.

Motif transitions were non-random on short timescales. For consecutive gaze shifts occurring within 500 ms, we computed transition probabilities between motifs and normalized each transition by the expected probability given the marginal fre-quency of the target motif (Methods 4.4). These transitions were organized around two patterns (Fig. 2E top): HD events preferentially transitioned back to HD, forming a recurrent head-dominant mode. By contrast, HwE, HbE, and ED formed a directed transition chain: HwE events were enriched for transitions to HbE, and HbE events were enriched for transitions to ED.

Direction-resolved transitions revealed that these two patterns had different relationships to movement direction (Fig. 2E bottom). HD provided the dominant transitions between nasal and temporal gaze shifts, consistent with repeated head-dominant sampling across directions, i.e., horizontal back-and-forth motion of the head. The HwE–HbE–ED chain was preserved within each movement direction. This transition structure was qualitatively stable across transition windows (Fig. S5), and the full direction-resolved observed-to-expected transition matrices are shown in Fig. S6. This is consistent with a sequence of active exploration initiated by a joint head-eye orienting movement (HwE), followed by continued head-led gaze shifts during movement (HbE), and a final eye-dominant movements at the end (ED).

Together, these analyses show that the four motifs are not merely local subdivisions of head–eye velocity space. They retain distinct timing signatures, occur in different locomotor speeds, and participate in organized short-timescale transitions.

### 2.3 Pre-onset visual-behavioral structure carries motif-specific information

We next tested whether the motifs also differed in the visual-behavioral input that preceded movement onset. Because the world-camera view is generated by the animal exploring, and the environment contains both static and dynamic visual stimuli, pre-onset video does not isolate external visual input. Rather, it reflects a joint product of the surrounding scene and ongoing self-motion. We therefore treated the pre-onset world-camera signal as a joint visual-behavioral measure and asked whether its structure carried information about which motif occurs next, rather than whether vision *per se* predicts the upcoming movement.

We analyzed retinocentrically corrected world-camera video in the window from *−*600 ms to *−*100 ms relative to gaze-shift onset (Fig. 3A; Methods 4.5). This window captured recent eye-centered input while excluding the peri-onset interval, when head and eye motion could dominate measured image change. For each event, we first quantified changes in local image contrast across the pre-onset window. Contrast change differed across motifs (Fig. 3B). HwE and ED events were preceded by significant increases in contrast, whereas HbE and HD events showed no significant positive contrast change. Thus, increasing pre-onset image contrast was associated with a subset of motifs rather than with gaze-shift onset generally.

**Fig. 3.**
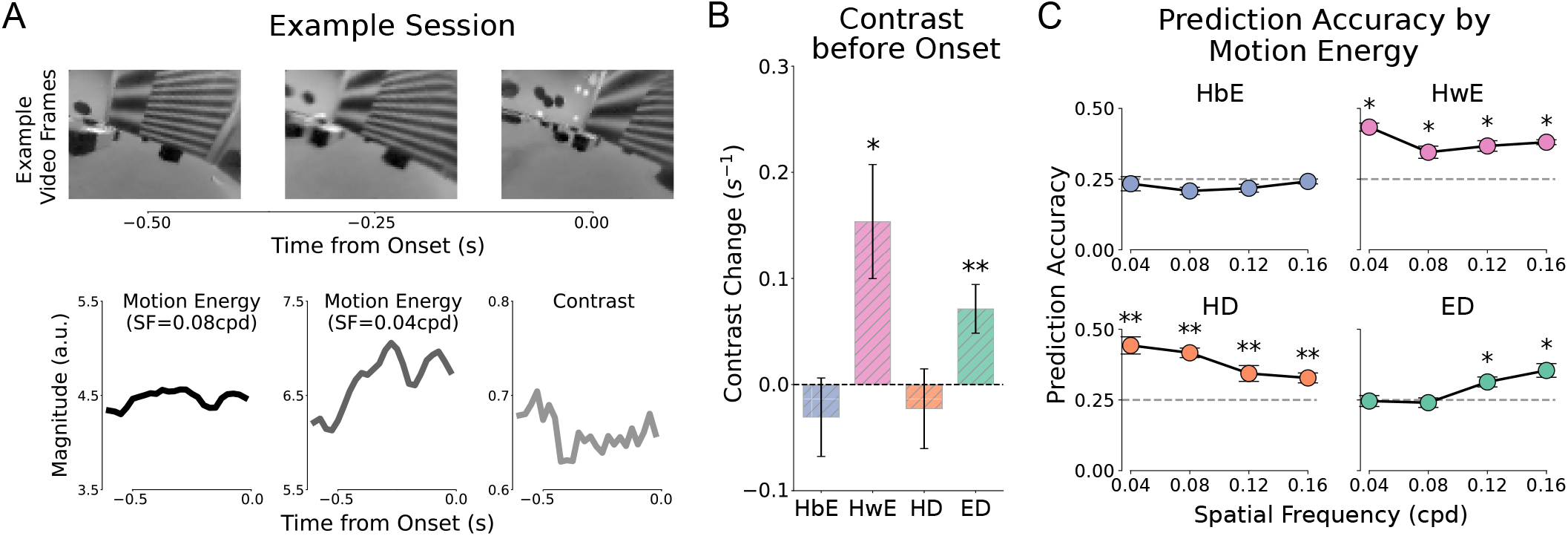
Pre-onset video features predict gaze-shift motif identity. **A,** Example world-camera frames preceding gaze-shift onset, shown with corresponding pre-onset video feature traces. Motion energy was extracted at multiple spatial frequencies; example traces are shown for 0.08 and 0.04 cycles per degree (cpd), together with image contrast. Video features were analyzed in the pre-onset window to test whether scene structure predicted subsequent gaze-shift motifs. **B,** Mean contrast change in the pre-onset window from *−*600 ms to *−*100 ms relative to gaze-shift onset across events, shown separately for each motif. Bars show mean *±* s.e.m. Asterisks indicate significant deviation from zero using a Wilcoxon signed-rank test with FDR correction. **C,** Motif-specific prediction accuracy from pre-onset motion energy. Support vector machine classifiers were trained to predict the gaze-shift motif from motion-energy features computed in the pre-onset window from *−* 600 ms to *−*100 ms relative to gaze-shift onset. Features were computed across spatial-frequency bands (0.04, 0.08, 0.12, and 0.16 cpd). Accuracy is shown as a function of spatial frequency after averaging across temporal-frequency bands. Error bars show mean *±* s.e.m. across cross-validation folds. Dashed line indicates chance performance (0.25). Asterisks indicate above-chance prediction after FDR correction (\**p* <0.05, \*\**p* <0.01).

Richer spatiotemporal features of the pre-onset signal also differed by motif. We computed direction-averaged motion energy from eye-centered pre-onset video frames across spatial- and temporal-frequency bands, then trained class-balanced support-vector-machine classifiers to read out the upcoming motif from these features (Fig. 3C; Methods 4.5). Read-out accuracy was assessed separately for each motif (chance = 0.25). Time-resolved results split by temporal-frequency band are shown in Fig. S7, and the full time-scale analysis in Fig. S8.

Here we summarized accuracy over the *−* 600 to *−*100 ms window as a function of spatial frequency, averaged across temporal-frequency bands. HbE events were not read out above chance at any spatial frequency. In contrast, HwE and HD events were distinguished from low-spatial-frequency motion energy, whereas ED events were distinguished primarily from higher spatial-frequency bands. The pre-onset visual-behavioral signal thus carried motif-specific information that was not uniform across motifs and depended on spatial scale.

We emphasize that these features cannot separate an external visual contribution from the visual consequences of self-motion. However, shifting event times by 10 s reduced read-out to chance (Fig. S8B), indicating that prediction is event-locked. In addition, matching the average body and gaze speed across motifs, and separately matching the optic-flow magnitude time-course across motifs over the same window, resulted in read-out broadly consistent with the full-data accuracy (Fig. S8C). These speed- and flow-matched controls indicate that the motif-specific information is not fully explained by body-gaze kinematics or by the pre-shift optic flow traces.

These analyses show that pre-shift structure carries information about the upcoming motif beyond optic-flow magnitude and body-gaze-speed signals. HbE showed little such pre-shift structure, whereas HwE, HD, and ED were distinguished by different components. For HwE, this signature converged with its behavioral profile identified above: near-synchronous head–eye onset, post-onset acceleration of body speed, and position at the start of the direction-preserving HwE–HbE–ED sequence. Whether any component of this pre-onset structure reflects specific visual input that helps drive the movement will require causal manipulations in future work.

### 2.4 Motif-specific gaze-shift responses are strongest and earliest in sSC for HwE

We next asked whether these motifs were also accompanied by distinct neural responses. Previous work has shown that responses in mouse primary visual cortex (V1) and superior colliculus (SC) are shaped by behavioral state (Franke et al., 2022; Ito et al., 2017; Savier et al., 2019), and that both V1 and superficial SC (sSC) are modulated around gaze shifts in freely moving mice (Parker et al., 2023; Sharp et al., 2025).

We recorded single-unit activity from V1 and sSC while simultaneously tracking head and eye movements in freely moving mice (Methods 4.6). Neural responses were aligned to gaze-shift onset and averaged across events for each motif. Both areas showed peri-onset modulation, but the motif dependence differed across regions (Fig. 4A). V1 responses were dominated by broadly consistent post-onset responses, whereas sSC responses varied more strongly across motifs around gaze-shift onset.

**Fig. 4.**
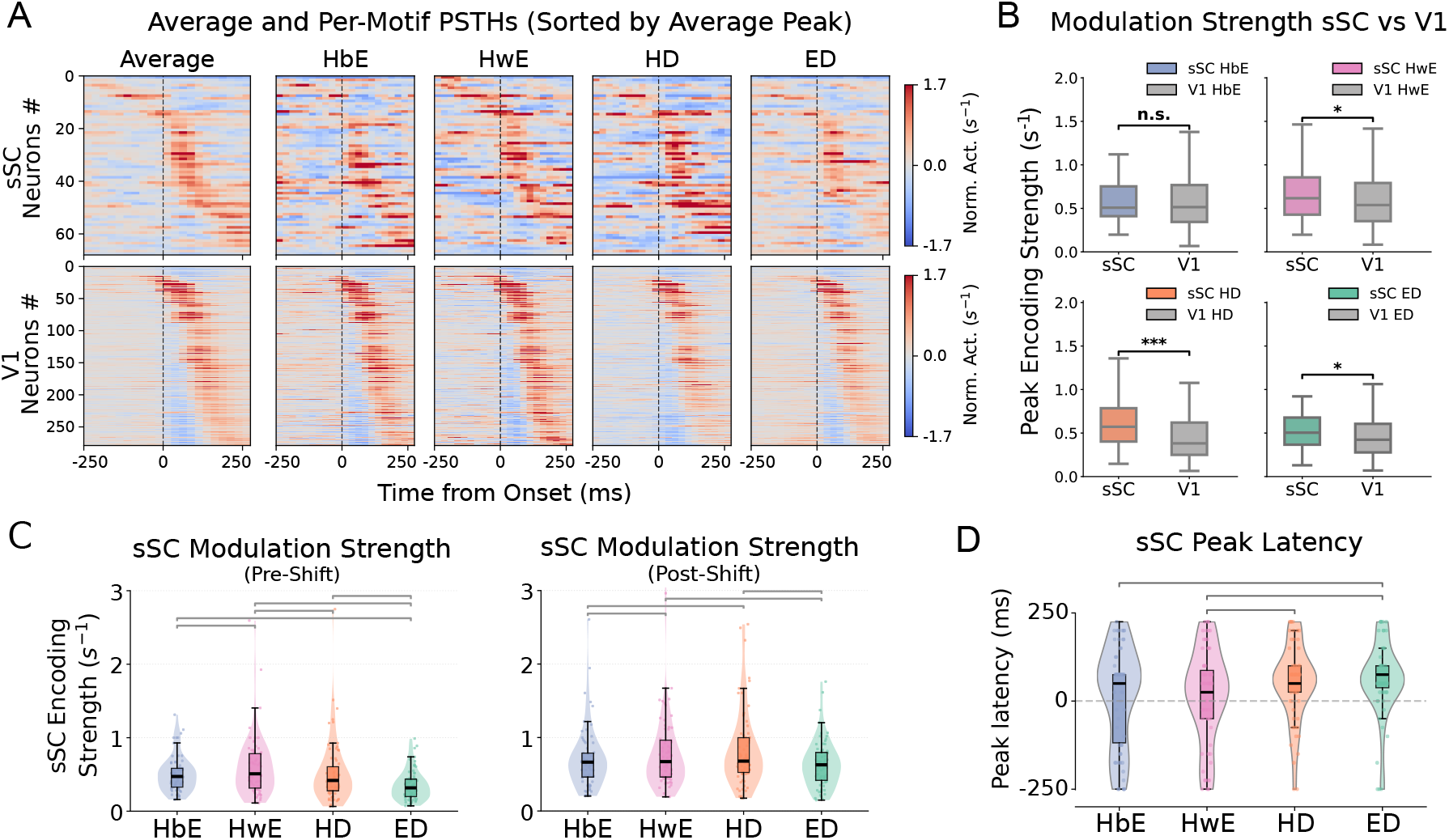
sSC responses are stronger and earlier for HwE. **A,** Averaged and motif-specific neural activity aligned to gaze-shift onset in superficial superior colliculus (sSC; left) and primary visual cortex (V1; right). Each row represents one neuron, sorted by peak response latency in the averaged response. Activity was baseline-subtracted and normalized within each neuron; color indicates normalized firing-rate modulation relative to the pre-event baseline. **B,** Peak fractional modulation within *±* 250 ms of gaze-shift onset, compared between sSC and V1 for each motif. Peri-event modulation strength was defined as the peak absolute baseline-normalized response, *|* (*R*_peak_ *−R*_baseline_)*/R*_baseline_ *|*, within the peri-onset window. Boxes show median and interquartile range (IQR); whiskers extend to 1.5 *×* IQR. **C,** Peak peri-event modulation strength across motifs within sSC, computed separately before gaze-shift onset (*™* 250 ms to 0; left) and after onset (0 to +250 ms; right). Violins show kernel-density estimates; boxes show median and IQR. Horizontal brackets indicate significant pairwise comparisons after FDR correction. **D,** Peak response latency within *±* 250 ms of gaze-shift onset, compared across motifs within sSC. Dashed horizontal line marks gaze-shift onset. Horizontal gray brackets indicate significant pairwise comparisons after FDR correction. *^∗^p <* 0.05, *^∗*^p <* 0.01, *^∗**^p <* 0.001; n.s., not significant; FDR-corrected tests as described in Methods. sSC, *n* = 68 neurons; V1, *n* = 279 neurons.

We quantified peri-event modulation strength as the peak ab-solute fractional modulation within *±* 250 ms of gaze-shift onset. Across areas, sSC showed stronger modulation than V1 for HwE, HD, and ED, but not for HbE (Fig. 4B). This regional difference aligned with the visual-behavioral analysis: the three motifs with stronger sSC modulation were also predicted above chance from pre-shift motion energy, whereas HbE, the least predictable motif, showed no significant sSC–V1 difference.

We then compared motif-dependent modulation within sSC before and after gaze-shift onset (Fig. 4C). In the pre-onset window, HwE produced the strongest sSC modulation among the four motifs. In the post-onset window, HD and HwE produced the strongest sSC modulation, indicating that motif-dependent sSC responses persisted after movement onset.

Finally, response timing differed across motifs in sSC. HwE responses peaked significantly earlier than HD and ED responses (Fig. 4D), indicating that motif-dependent sSC responses differed in both magnitude and timing. Together, these results show that gaze-shift motifs are accompanied by distinct peri-onset responses in V1 and sSC. Note that the neural data do not establish a causal role. Instead, they provide convergent evidence: the motifs most predictable from pre-onset visual-behavioral input showed stronger sSC than V1 modulation, and HwE combined the strongest pre-onset sSC modulation with the earliest sSC peak. This convergence supports HwE as a candidate active orienting motif in freely moving mice.

## 3 Discussion

Mouse gaze shifts during natural exploration are usually treated as a single event class. Our results show that this operational category is more diverse than its single-class treatment implies, revealing multiple reproducible head–eye coordination motifs with distinct behavioral and neural signatures. Using time-resolved peri-event kinematics, we identified four dominant motifs: Head-before-Eye (HbE), Head-with-Eye (HwE), Head-Dominant (HD), and Eye-Dominant (ED), that differed in head–eye timing, locomotor dynamics, short-timescale sequence structure, pre-onset visual-behavioral structure, and peri-event modulation in V1 and sSC. What appears as a uniform behavior under coarse instantaneous classification thus contains a structured repertoire of gaze-control modes.

Among these, HwE emerged as a candidate active gaze-shift mode not previously described in freely moving mice. HwE events showed near-synchronous head and eye onset, began from a more stationary head, were followed by increased body speed, were predictable from pre-shift visual-behavioral features, and elicited the strongest pre-onset and earliest peri-onset sSC modulation. A near-synchronous head–eye onset that initiates a movement is a signature of active orienting rather than image stabilization. This profile resembles the stimulus-driven, SC-dependent saccades described in head-fixed mice by Zahler et al. (2021) and the tightly coupled eye-head co-initiated gaze shifts reported in body-fixed mice by Verdone et al. (2026). HwE extends these observations to freely moving behavior, identifying a combined head–eye orienting movement during natural exploration. Because our measures are correlational, causal manipulation of external stimuli or SC activity will be required to determine whether either is necessary to generate the motif in freely moving mice.

The pre-onset sSC signal is particularly informative for this interpretation. Previous work has emphasized peri-gaze responses in V1 and SC as consequences of gaze shifts (Parker et al., 2023 ; Sharp et al., 2025). Here, motif-specific sSC modulation was already present before movement onset and was strongest for HwE. This pre-onset modulation points to preparatory activity or a drive from the pre-onset scene, or both. Distinguishing these alternatives will require perturbation or closed-loop experiments, but the timing is in alignment with sSC responding to the initiation of HwE rather than only responding to its consequences.

The remaining motifs occupied different behavioral regimes. HD events were head-dominated and provided the dominant transitions between nasal and temporal gaze shifts. This profile is consistent with repeated head-based sampling of the scene during slower exploration (Wallace et al., 2013), in line with the broader reliance on head movements for gaze sampling in afoveated species (Land, 2019).

HwE, HbE, and ED formed a directed transition chain that, unlike the recurrent HD mode, was preserved within a single movement direction. The ordering suggests a stereotyped progression: an initial combined head–eye movement (HwE) commits gaze to a new direction, followed by head-led gaze shifts (HbE) while the head is still in motion, and finally eye-dominant movements (ED) as the head decelerates and finer adjustment is achieved primarily by the eye. The transition from HwE to HbE is consistent with an active orienting movement giving way to a “saccade and fixate”-like regime. HbE was head-led, weakly distinguished by pre-onset visual-behavioral structure, and showed no stronger sSC than V1 modulation. Consistent with a more reflexive role, the eye counter-rotated against the head before the gaze shift, and the saccade then reset eye position in the orbit, a pattern characteristic of the “saccade-and-fixate” strategy (Land and Tatler, 2009; Meyer et al., 2020; Michaiel et al., 2020). We note that this analysis is observational: motif identity predicts which motif follows, but does not establish that one motif mechanically generates the next.

Notably, because the HwE and HD motifs depend on the head being stationary either before or during the gaze shift, these types would be much less frequent during active behaviors such as prey capture (Michaiel et al., 2020) or tracking (Meyer et al., 2020) than during the spontaneous alternation between stationary periods and free exploration examined here. This may explain why these previous studies primarily observed the saccade-and-fixate pattern (i.e., HbE). It will be interesting to determine how the different motifs are engaged during goal-directed behavior: for example, a bout of prey pursuit might be predicted to begin with an HwE orienting movement.

Several limitations exist. First, the motifs are statistical modes of an underlying continuous kinematic space. The four-motif partition should be understood as a reproducible and useful decomposition, not as a strict categorical boundary between discrete behaviors. Second, all analyses are correlational. Causal tests could combine closed-loop manipulations with perturbations, for example using optogenetic approaches (Zahler et al., 2023), virtual-reality preparations for rodents (Aronov and Tank, 2014 ; Grosso et al., 2017), or mouse digitaltwin analysis (Xu et al., 2023 ; Lima et al., 2026). Third, V1 and sSC are only part of the circuit. Deep SC has been reported to be modulated by head movements during gaze shifts (Sharp et al., 2025), and our four motifs differed in head motion. The pulvinar carries saccade-related signals to V1 that help distinguish self-generated from external motion (Miura and Scanziani, 2022), and secondary motor cortex contributes to head–eye movements in mice (Itokazu et al., 2018 ; Sato et al., 2019 ; Barthas and Kwan, 2017). Future work should test whether these regions carry motif-specific signals and whether they help select among coordination modes.

These findings revise the prevailing view of mouse gaze control during natural behavior. Rather than being limited to image stabilization and reflexive head-coupled resetting, freely moving mouse gaze includes a structured repertoire of coordination motifs with distinct relationships to locomotion, visual-behavioral input, and neural activity. Among these, HwE suggests that an active head–eye orienting mode, which had previously been characterized only in head-fixed (Zahler et al., 2021) and body-fixed (Verdone et al., 2026) preparations, is also expressed during free movement. Our results establish that natural mouse gaze is more diverse and more active than a single reactive class, comprising distinct head–eye coordination modes with their own behavioral and neural signatures. By identifying active orienting within the behavioral repertoire, they support the view that active gaze control is a shared feature of mouse vision rather than a property of foveated animals or constrained preparations alone.

## 4 Methods

### 4.1 Animals, electrophysiology, and behavioral recordings

Data were collected by and described in detail in Parker et al. (2023) and Sharp et al. (2025). Here, we provide a brief summary.

All procedures were conducted in accordance with National Institutes of Health guidelines. 3-10 month-old mice (C57BL/6J, Jackson Laboratories and bred in-house) were kept on a 12-h light/dark cycle. Mice were housed with sibling cage-mates until the electrophysiology implant. Humidity was 40% - 60% and temperature was 21 *±* 1°C. In total, 11 mice (4 from primary visual cortex recordings and 7 from superior colliculus recordings) were used for this study. Mice of both sexes were pooled for analysis.

Freely moving mice were implanted with a titanium head-plate over either the superior colliculus (SC) or the primary visual cortex (V1), along with a miniature connector for reversible attachment of head-mounted hardware. After habituation, a linear silicon probe (Diagnostic Biochips, 64 or 128 channels) mounted in a custom 3D-printed microdrive was implanted through a craniotomy over the target region (1,500 *µm* below the pia for SC; 750 *µm* for V1), with a reference wire placed over the left frontal cortex. Recordings began the day after implantation. Neural signals were acquired at 30 kHz (bandpass 0.01 Hz-7.5 kHz) using an Open Ephys system. Head-mounted hardware consisted of an eye-facing camera with infrared illumination (30 fps), a forward-facing world camera (BETAFPV C01, 30 fps), and a three-axis IMU (Rosco Technologies, 30 kHz, downsampled to 300 Hz). A top-down camera (FLIR Blackfly, 60 fps) tracked the animal in the arena. Acquisition was performed in Bonsai using custom workflows (github.com/nielllab/FreelyMovingEphys), and all data streams were aligned offline via system timestamps. In each session, each animal was placed in a 48 *×* 37 *×* 30 cm arena for approximately one hour of free exploration. Three arena walls were covered with patterned wallpaper (gratings and white noise), and the fourth displayed a moving sparse noise stimulus; the floor was densely covered with black and white Lego bricks for 3D contrast, with small food pieces scattered to encourage foraging (animals were not food-or water-restricted).

### 4.2 Behavioral and neural data pre-processing

Full preprocessing details are described in Parker et al. (2023) and Sharp et al. (2025), and we summarize the relevant steps here.

For each session, raw electrophysiology data from freely moving recordings were collected. Common-mode noise was removed by subtracting the median across channels at each time point, and spikes were sorted with Kilosort 2.5 (github.com/MouseLand/Kilosort). Single units were then curated in Phy 2.0 (github.com/cortex-lab/phy) based on contamination (*<* 10%), mean firing rate (*>* 0.5 Hz across the full recording), waveform shape, and autocorrelogram structure. After curation, spike trains were split back into individual recordings. Laminar position was estimated from the multi-unit LFP recorded during head-fixed visual stimulation. For V1, the LFP on each channel was bandpass filtered (1-300 Hz), and the channel with maximum LFP power was taken as the center of layer 5. For SC, evoked LFP responses to a contrast-reversing checkerboard were averaged across reversals, and the time point of maximum negative amplitude was used to identify the center of the superficial SC (sSC). Units within 0-300 *µm* of the sSC center were classified as sSC.

The animal’s position in the arena was tracked from the top-down camera with DeepLabCut (Mathis et al., 2018), and running speeds were measured by the neck-point velocities. Eye camera video was deinterlaced to 60 fps, and eight points around the pupil were tracked with DeepLabCut; an ellipse was fit to these points, and pupil position was expressed as angular rotation. World camera video was deinterlaced to 60 fps and lens distortion corrected with OpenCV. Horizontal head rotation velocity was extracted from the IMU, converted to deg/s, and interpolated to eye camera timestamps, with leftward and rightward defined from the animal’s perspective (i.e., nasal motion corresponds to a leftward movement, and temporal motion corresponds to a rightward movement).

### 4.3 Gaze shift detection and selection

To identify gaze shift events, we used horizontal head rotation velocity (extracted from the IMU) and horizontal eye velocity (computed as the temporal derivative of horizontal pupil angular position). Eye velocity and head velocity were aligned to a common 25-ms time base and were both smoothed with a Savitzky-Golay filter. We defined gaze velocity as head plus eye velocity at each time point. We then performed an initial classification of each time point into nasal gaze shifts, temporal gaze shifts, or compensatory movements. The joint (head velocity, eye velocity) features were projected into a 4-dimensional space using Gaussian random projection and clustered with k-means (k = 3). The three resulting clusters were assigned to nasal gaze shifts, temporal gaze shifts, and compensatory movements based on the sign of each cluster’s mean gaze velocity. Because SC recordings were noisier than V1 recordings, a small fraction of SC events whose head and eye velocities had clearly opposing signs were reassigned to the compensatory class. Classification was performed separately for each region (sSC, V1)*×* direction (nasal, temporal) combination using otherwise the same pre-processing pipeline. Gaze-shift-classified time points whose immediately preceding frame was labeled as compensatory were excluded.

To further partition gaze shifts into functional motifs based on their temporal dynamics, we constructed a temporal feature vector for each event. At each event onset, we sampled horizontal head velocity, eye velocity, head acceleration, and eye acceleration (acceleration computed as the time derivative of the smoothed velocity) at *±* 100 ms surrounding each gaze shift event. Features were z-scored and clustered with k-means (k = 4). To minimize contamination from temporally adjacent events of different motifs, any gaze shift whose neighboring frames within 100 ms were assigned to a different cluster was excluded. For each remaining run of consecutive same-motif-category frames, the first data point was then taken as the event onset.

To ensure each motif was adequately sampled, we required at least ten events per motif and gaze shift direction; mice below this threshold in any of the four motifs were excluded from all downstream analyses. This criterion excluded three SC mice and zero V1 mice, leaving n = 4 mice per region for the analyses reported below. The resulting data were carried forward as the functional gaze-shift motifs for all subsequent analyses.

### 4.4 Behavioral context

Events from both directions (nasal and temporal) and both regions (sSC, V1) were pooled by motif label and sorted in time within each recording unless otherwise stated. In head-eye difference and onset lag analyses, to allow pooling across directions, temporal (right shift) events were sign-inverted so that head and eye velocities were expressed in a common nasal-direction reference frame. For each event, we extracted a *±* 100 ms window around each event onset. For this analysis, we defined the onset time of head and eye motion separately as the first frame in the window at which the trace exceeded 50% of its peak value. The head-eye lag was then computed as head’s onset - eye’s onset, in milliseconds, such that negative lags indicate that the head began moving before the eye. Events in which only the eye or only the head moved were assigned to placeholders so that their relative frequency was retained in the cumulative distributions.

To test whether specific gaze-shift motifs preferentially followed others, we constructed a transition matrix between consecutive events. For each event, we identified the next event of any motif occurring within the same recording; only consecutive pairs separated by *≤*500 ms were retained as valid transitions to measure the gaze shift sequence within a short interval. The observed transition matrix *P* (*j| i*) was computed by counting transitions from motif *i* to motif *j* across all valid pairs. To express each transition relative to chance, we computed a lift matrix by dividing the observed probability by the marginal frequency of the target motif. Values *>* 1 in our lift matrix indicate that the transition occurred more often than expected from the motifs’ overall frequencies. Statistical significance was measured via a within-mouse label permutation test. For each of 10,000 permutations, gaze shift labels were randomly shuffled within each mouse while preserving the original inter-event intervals and the set of valid transition pairs. The transition count matrix was recomputed on each shuffle to generate a null distribution for every cell. *z*-scores were obtained by comparing each observed count to its null distribution, and transitions were considered significantly enriched at *z >* 1.96.

### 4.5 Video features

To measure whether different gaze-shift motifs were preceded by distinct video features, we analyzed the head-mounted world camera video with retinocentric correction, which shifted each frame for the animal’s current eye position so that the analyzed image approximates the scene on the retina (see Parker et al. (2022b) for details). For each gaze shift event, we extracted a pre-event window spanning *−*600 to *−*100 ms relative to event onset. This time window was chosen to capture the more immediate scene the animal experienced before gaze shift while controlling the change from the head movements during the gaze shift itself. Within each frame, we computed local Michelson contrast at every pixel as (*I_max_− I_min_*)*/*(*I_max_* + *I_min_* + *ϵ*), where *ϵ* = 10*^−8^*. To quantify how rapidly contrast changed, we fit a sliding linear regression to the contrast time series within the pre-event window. Slopes were computed over consecutive 100-ms windows, and the peak rate of contrast change for each event was defined as the slope with the largest absolute value across all windows. This per-event value (in units of contrast per second) was used for statistical comparison. For each motif, we tested whether the rate of contrast change differed significantly from zero using a Wilcoxon signed-rank test.

To extract direction-agnostic spatiotemporal motion energy from the videos, we first filled the missing-pixel periphery caused by eye displacement in retinocentric frames. We then used a Gabor motion-energy pyramid (moten package, Nishi-moto et al. (2011)) with a filter temporal width of 10 frames. Pyramids were constructed for each combination of multiple spatial frequencies (0.04-0.16 cpd) and temporal frequencies (4-16 Hz). For each SF-TF pair, motion energy was averaged across the eight orientation channels of the pyramid to yield a direction-invariant feature. For each gaze shift event, we extracted a *±* 1 s peri-event window (81 frames at 40 Hz sampling rate) of motion energy from each event onset.

To test whether video features predicted gaze-shift motifs, we trained classifiers to discriminate the four motifs from motion-energy features at each SF-TF-time combination. Nasal and temporal events were pooled. The 81-bin peri-event motion energy was partitioned into 20 non-overlapping windows of 4 frames (100 ms) each, spanning approximately -1 s to +1 s. For each SF-TF-time, we performed B bootstrap iterations, where B equals the largest motif’s event count; on each iteration we sampled N events without replacement from each motif, where N equals the smallest motif’s event count. This B and N subsampling enforced class balance while utilizing all data points. Samples were split 70/30 into train/test sets; features were *z*-scored; and a multi-class support vector machine was fit to the training set and evaluated on the held-out test set. Per-class accuracy was averaged across bootstrap iterations. To summarize motif-specific read-out as a function of spatial and temporal frequency, we focused on a pre-event time window that covers video information from -600 to -100 ms before onset, after taking the temporal filter window width into account. For each motif and each SF-TF, we took the maximum per-class accuracy across the pre-event time windows. SF performance curves were obtained by averaging across TFs, and TF performance curves by averaging across SFs. For each frequency, accuracy was tested against chance (0.25) using a one-sample Wilcoxon signed-rank test, and p-values were corrected across frequencies using the Benjamini-Hochberg false discovery rate procedure (*α* = 0.05).

### 4.6 Neural response

Single-unit recordings were obtained from the superficial superior colliculus (sSC; n = 4 mice) and primary visual cortex (V1; n = 4 mice). For each gaze shift event, the central *±* 500 ms around onset were extracted as the peri-event response, and*−*750 to *−*500 ms relative to onset served as the pre-event neural response baseline. Events of the same mouse and motif were averaged.

For each mouse-neuron pair, baseline samples were concatenated across all four motifs to form a single global baseline. Each motif-specific peri-event trace was then zero-centered by the global baseline mean and divided by its absolute value, yielding a fractional change from baseline. Using this common baseline across motifs preserves within-neuron differences across gaze-shift motifs.

Peri-event modulation strength was defined as the peak absolute fractional modulation in each motif and each neuron within a fixed *−*250 ms to +250 ms response window around event onset. Peak latency was defined as the time of this peak relative to event onset. To restrict latency analyses to neurons that were meaningfully modulated by gaze shifts, neuron– motif pairs whose peak amplitude fell below 0.25 were excluded. Cross-region comparisons within each motif (sSC vs. V1) used the Mann-Whitney U test for peri-event modulation strength and peak latency. Within-region comparisons across motif pairs used the Wilcoxon signed-rank test on paired per-neuron values.

### 4.7 Statistics

All analyses were done using Python (v 3.9). The error bars in the figures represented the standard error of the mean unless otherwise stated. We used the Wilcoxon signed-rank test, Mann-Whitney U test, and permutation tests for statistical significance. All statistical analyses were done with Benjamini-Hochberg FDR correction wherever applicable. The significance level was 0.05. Significance thresholds are reported as * p *<* 0.05, ** p *<* 0.01, *** p *<* 0.001.

### 4.8 Data and code availability

Data were publicly available at Parker et al. (2023) and Sharp et al. (2025). All code will be publicly available upon publication.

## A Motif discovery and clustering robustness

**Fig. S1.**
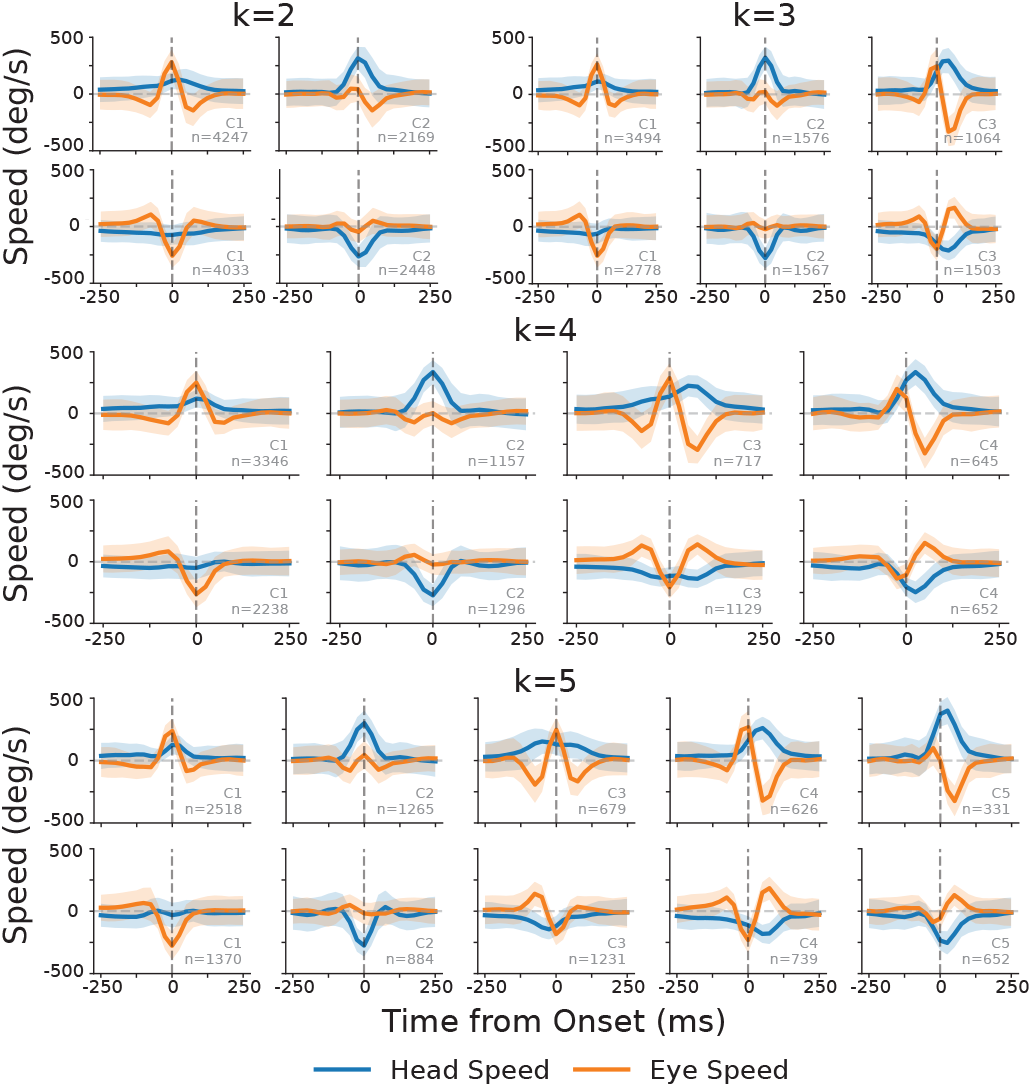
Motif discovery using different numbers of ks.

**Fig. S2.**
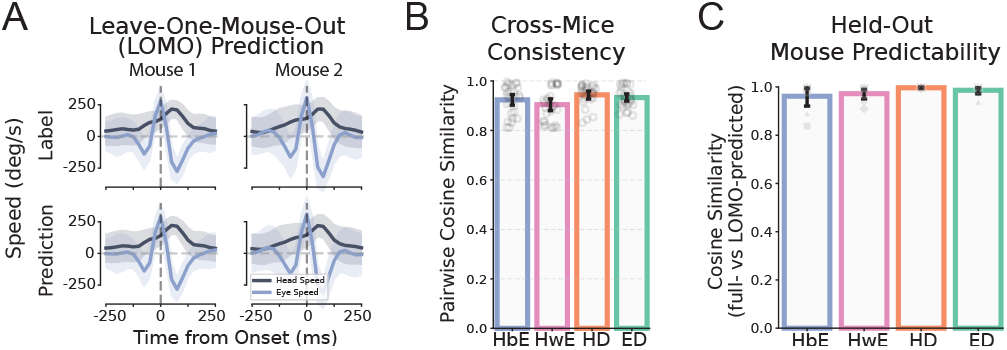
Leave one mouse out cross-validation for clustering consistency. **A,** Example head-eye movement trajectories using full data k-means clustering (**top**) and using leave-one-mouse-out LOMO to predict the held out mouse (**bottom**). **B,** Pair-wise cross-mice consistency in LOMO prediction; **C,** Held-out mouse predictability in LOMO.

To determine the appropriate number of partitions, we balanced two considerations: identifying the biologically distinct gaze-shift dynamics and maintaining reliability across animals. At k=2 and k=3 (Fig. S1), the partition failed to separate Head before Eye and Head with Eye movements, which have been characterized as functionally distinct motifs in head-fixed experiments (Zahler et al., 2021) and in body-fixed experiments (Verdone et al., 2026). At k=4, this distinction emerged. Cross-mouse consistency and leave-one-mouse-out predictability were reduced relative to k=3 (Fig. S2), but the scores remained high (consistency: 0.963 *±* 0.029 at k=3 vs. 0.926 *±* 0.055 at k=4, Wilcoxon signed-rank *p* <0.001; pre-dictability: 0.996 *±* 0.005 vs. 0.979 *±* 0.037, *p* <0.001). In-creasing to k=5 further reduced consistency and predictability (0.883 *±* 0.092, *p* <0.001; and 0.949 *±* 0.064, *p* <0.001, re-spectively) without introducing a qualitatively new dynamic: C4 and C5 at k=5 corresponded to a subdivision of C4 at k=4 along a post-shift temporal axis rather than a categorical boundary documented in prior work. In other words, the reliability cost from k=3 to k=4 provides a literature-anchored distinction; the comparable cost from k=4 to k=5 buys only a finer slice within an existing dynamics class. We therefore picked k=4 as a partition that captures the established categorical distinctions while preserving cross-animal reliability.

All values are mean *±* SD across mouse-pairs (consistency) or held-out mice (predictability). The leave-one-mouse-out (LOMO) validation was done by fitting the k-means classifier using all but one mouse’s data, then predicting the label in the left-out mouse’s data. The predictability score was measured by the cosine similarity between the predicted motif label from LOMO and the actual motif label from using the full-data classifier; the cross-mouse consistency was measured by the average pairwise cosine similarity of the predicted labels across mice.

## B Temporal/nasal pooling

**Fig. S3.**
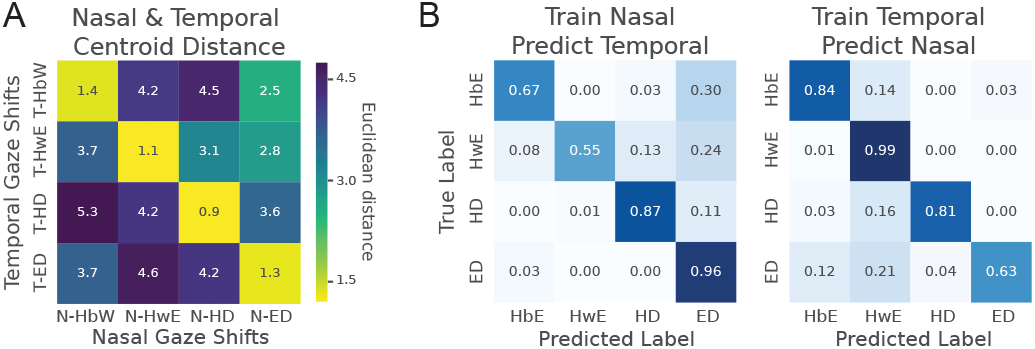
Nasal/Temporal gaze shift matching. **A,** Euclidean distance matrix between per-motif centroids in nasal and temporal computed in trajectory feature space. Matched pairs (di-agonal) are 2.2 *×* - 4.5 *×* smaller than unmatched pairs. **B,** Confusion matrices for cross-direction classification. **Left**: classi-fier trained on nasal events and tested on temporal events (accuracy 77%). **Right**: classifier trained on temporal and tested on nasal (accuracy 78%). Within-direction 5-fold cross-validation accuracies are 94% (nasal) and 93% (temporal); chance is 25%.

We examined whether the four motifs are recovered across movement directions. To isolate trajectory shape, we signflipped temporal traces and z-scored each kinematic input variable (head speed, eye speed, head acceleration, eye acceleration) separately per direction before comparing.

We first computed the per-motif mean centroid in each direction and the 4 *×*4 Euclidean distance matrix between nasal and temporal centroids (Fig. S3A). For every motif, the near-est centroid across directions was the matched counterpart, and the Hungarian optimal one-to-one assignment was the identity. Diagonal (matched-pair) distances were 2.2*×* to *×* 4.5 smaller than the mean off-diagonal distance within the same row.

We then trained a multinomial logistic regression classifier on the trajectory features of one direction (nasal or tempo-ral) and tested it on held-out events from the current direction (nasal or temporal) or the other direction (temporal or nasal). See Fig. S3B. Within-direction cross-validated accuracy was 94.53% (train and test on nasal) and 92.52% (train and test on temporal); cross-direction accuracy was 77.47% (train on nasal, test on temporal) and 77.98% (train on temporal, test on nasal), well above the 25% chance.

In summary, these results indicate that the four-motif structure is preserved across gaze-shift directions, justifying pooling within each motif in subsequent analyses.

**Fig. S4.**
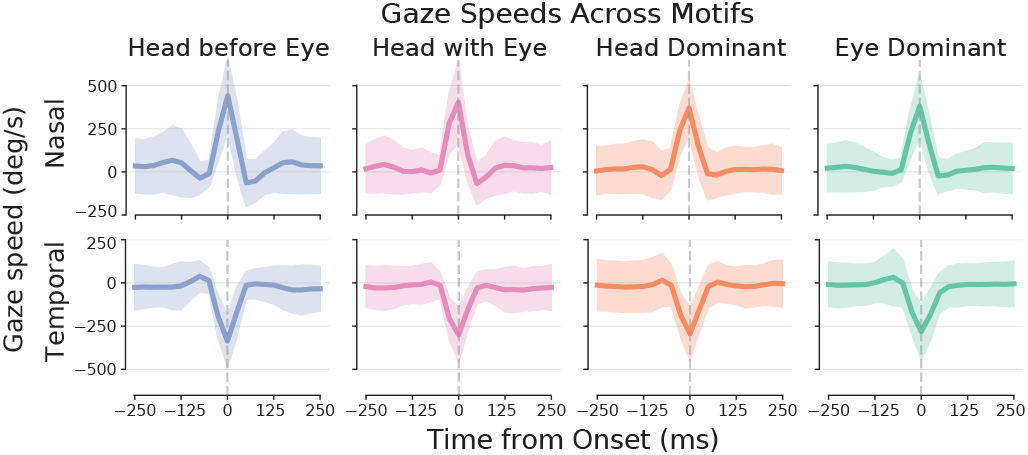
Gaze speed profiles *±* 250 ms surrounding the shift onset for nasal (**top**) and temporal (**bottom**) movement direc-tions. Mean gaze speed (deg/s) *±* SD is shown across all events in each motif.

## C Transition probability by time and direction

**Fig. S5.**
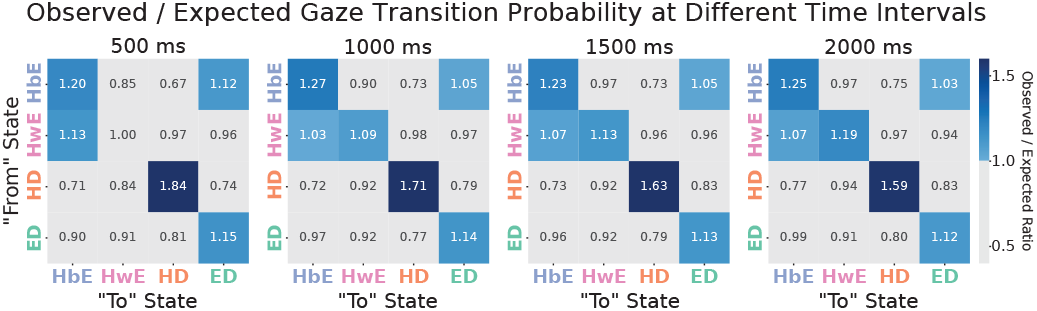
Sequence transition probability using different time intervals. Short-timescale transition structure among gaze-shift motifs. The matrix shows observed transition probability within 500 ms, 1000 ms, 1500 ms, or 2000 ms of each gaze shift, normalized by the expected probability given the overall frequency of the target motif. Values greater than 1 indicate preferred transitions and are colored in blue gradients.

## D Additional prediction analysis

**Fig. S6.**
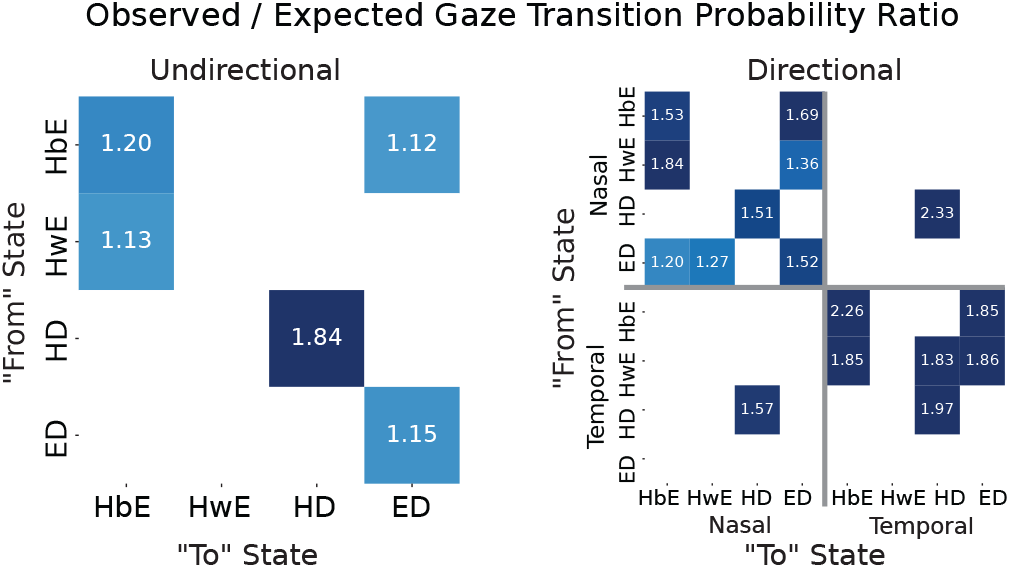
Direction-resolved transition matrices underlying Fig. 2E. Transition structure among gaze-shift motifs, with data either pooled across movement directions (left) or separated by nasal-to-temporal direction (right). The matrix shows the observed transition probability for consecutive gaze shifts occurring within 500 ms of each other, normalized by the expected probability given the overall frequency of the target state. States are defined by motif and movement direction. Values greater than 1 indicate transitions are preferred beyond chance expectation. Only significant transitions are shown (permutation test with > 95% confidence interval) and are colored in blue gradients.

**Fig. S7.**
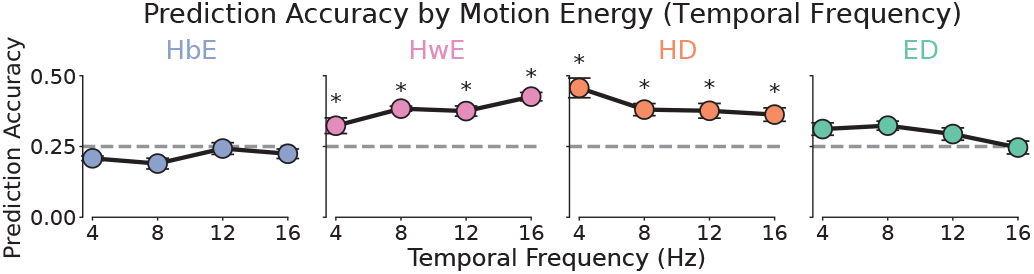
Motif-specific prediction accuracy from pre-onset motion energy. Support vector machine classifiers were trained to predict the gaze-shift motif from motion-energy features com-puted in the pre-onset window from *−*600 ms to *−*100 ms relative to the gaze-shift onset. Features were computed across temporal-frequency bands (4, 8, 12, and 16 Hz) after averaging across spatial-frequency bands. Error bars show mean *±* s.e.m.

In the main text, we showed that the spatial-frequency content of the pre-shift visual-behavioral input is predictive of the up-coming gaze-shift motif and reported per-SF prediction scores within the response window (averaged across temporal frequencies) as dot plots. Here we provide the per-TF prediction analysis (averaged across spatial frequencies) as dot plots in Fig. S7, as well as the full time-resolved version of that analysis, separately for motion energy computed from each SF and TF band (Fig. S8A) and a shifted-time null control (Fig. S8B).

As shown in Fig. S7, HwE and HD labels could be read out from motion energy generated by different temporal frequencies (HwE from relatively higher temporal frequency motion energy, and HD from relatively lower temporal frequency motion energy).

In terms of the time-shifted control, for every gaze-shift event, we computed local motion energy in a *±* 1 s window around shift onset following the procedure described in Meth-ods 4.5, and we trained an SVM to decode the four gaze-shift motifs (HbE, HwE, HD, ED) from the resulting motion-energy features. The 81-bin window was partitioned into 20 non-overlapping 100 ms bins (4 frames at 40 Hz), and a classifier was trained independently for each bin, yielding a prediction trace from -1 to +1 second (Fig. S8A). Panel A shows these traces broken down by SF bands (0.04, 0.08, 0.12, 0.16 cpd) averaging across TFs, and by TF bands (4, 8, 12, 16 Hz) averaging across SFs. The vertical dashed lines approximate the response window used for the dot plot (Fig. 3) in the main text. The horizontal dashed line indicates the four-way chance level (0.25). The prediction in motifs yielded above-chance performance across different time windows.

**Fig. S8.**
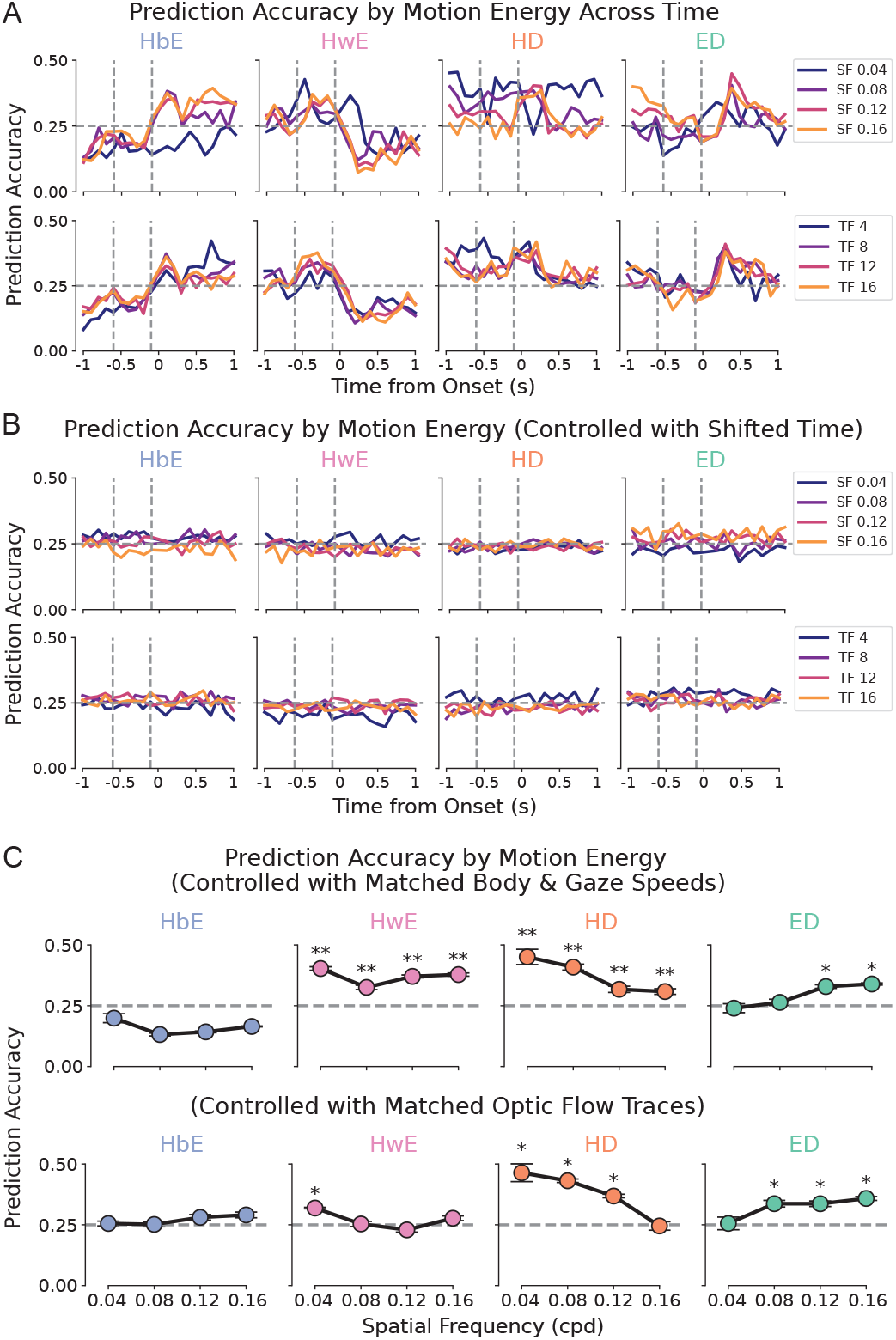
**A,** Prediction accuracy by pre-shift motion energy across time, where the motion energy is computed across spatial-frequency bands (0.04, 0.08, 0.12, and 0.16 cpd) and temporal-frequency bands (4, 8, 12, and 16 Hz). **B,** Same pre-diction pipeline but with shifted time as a control. **C,** Motif-specific prediction accuracy from pre-onset motion energy. Using matched body and gaze speeds (**top**) and with matched optic flow traces (**bottom**) as controls.

To verify that the above-chance decoding is not an artifact of session-level statistics shared across gaze events of the same motif, we repeated the entire pipeline on temporally displaced events. For each true gaze-shift event, we shifted the event time by *−*10 s, which keeps the sample within the same recording session (and therefore matches stimulus statistics, mouse identity, and recording state) while breaking its temporal relationship to the upcoming gaze shift (Fig. S8B Motion-energy features were extracted around these shifted timestamps, and the same SVM was trained in the same way as in panel A; the only methodological difference beyond the time shift is that for each classifier we drew a random set of 4 frames from within the 1*±* s window. Across all SF and TF bands and all four gaze-shift motifs, the controlled prediction accuracy fluctuates around the 0.25 chance line and never approaches the values observed inside the response window of panel A, confirming that the decodable information identified in the main text is specific to the short interval immediately preceding the gaze shift.

To confirm that the motif-specific motion energy decoding reflects differences that could not be entirely explained by pre-shift self-motion or retinal flow, we repeated the decoding after matching the four motifs on pre-shift covariates with two controls: First, in the speed control, we matched average body speed and average gaze speed (head plus eye velocity) between -600 and -100 ms from onset. Each variable was partitioned into 7 quantile bins, and within every joint bin, we subsampled an equal number of data points from each of the four motifs, so that the pooled set was balanced on the joint distribution of body and gaze speed across motifs. Second, in the optic-flow control, we matched on the full pre-shift traces of retinal optic-flow magnitude from -600 to -100 ms before onset (40 Hz sampling rate). Taking HwE as the reference, we paired each HwE event with its nearest neighbor from each of the other three motifs in flow-trace space (*k*-d tree, Euclidean distance) and retained only quadruplets in which all three neighbors fell within a thresholded distance, yielding events with near-identical pre-shift optic flow traces between -600 and -100 ms before onset. In both controls, the matched motion-energy features were passed to the same SVM as in panel A (Fig. S8C). The band-specific decoding performance reported in the main text largely persisted under both controls, indicating that it is not explained solely by the motif differences in pre-shift average gaze and body speed or pre-shift optic flow traces. Note that matching by subsampling distorts the within-motif data structure. For example, the matched optic-flow traces fall well below any motif’s original mean flow. Even with speed and flow controlled, the detailed pattern of movement within the window is not, so a residual contribution of self-motion to the prediction cannot be entirely excluded.

## E Neural analysis in V1

**Fig. S9.**
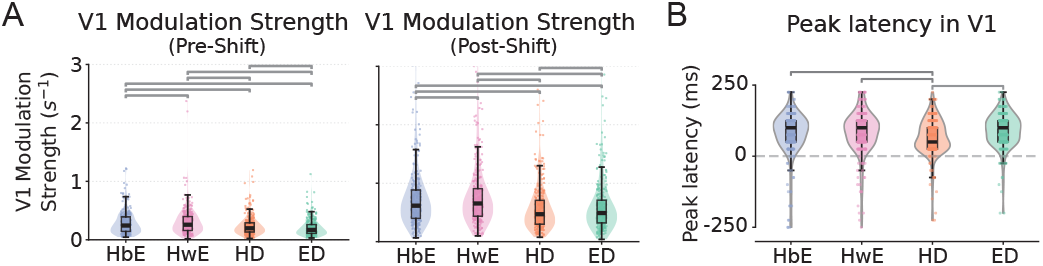
**A,**V1 modulation strengths pre- and post-gaze shift. **B,** V1 Peak latency relative to gaze-shift onset.

We performed the same analysis for motif modulation within V1, where the modulation is defined as the peak absolute fractional firing magnitude within the response window. During the pre-gaze-shift window, HwE produced the strongest modulation, significantly surpassing all other motifs (*p* <0.001 for all comparisons). HbE and HD exhibited intermediate modu-lation strengths that did not differ significantly from one another (p = 0.285), but both were significantly stronger than ED (*p* <0.001 for both). For post-gaze-shift responses, HwE again showed the strongest modulation, compared with HbE, ED, and HD (*p* <0.001 in all comparisons).

When assessing peak response latency in V1, HD responses peaked significantly earlier than ED, HbE, and HwE responses (*p* < 0.005 in all comparisons). There were no significant latency differences among the remaining three motifs (*p* > 0.05 for all comparisons).

## Footnotes

1 Because eye movements were recorded from a single eye, disconjugate components accompanying head pitch may appear as eye-only movements and contribute to a subset of ED events. Non-yaw head rotations can project onto the horizontal head-speed estimate under imperfect inertial-sensor alignment and contribute to a subset of HD events, though manual comparison against top-down tracking suggested these are relatively few. We retain both motifs but note these alternative contributions. A more controlled setup could test these coordination motifs more directly.

## References

Ambrad Giovannetti, E. and Rancz, E. (2024). Behind mouse eyes: The function and control of eye movements in mice. Neuroscience & Biobehavioral Reviews, 161:105671.

Aronov, D. and Tank, D. W. (2014). Engagement of Neural Circuits Underlying 2D Spatial Navigation in a Rodent Virtual Reality System. Neuron, 84(2):442–456.

Barthas, F. and Kwan, A. C. (2017). Secondary motor cortex: Where ‘sensory’ meets ‘motor’ in the rodent frontal cortex. Trends in Neurosciences, 40(3):181–193.

Dowell, C. K., Lau, J. Y. N., Antinucci, P., and Bianco, H. (2024). Kinematically distinct saccades are used in a context-dependent manner by larval zebrafish. Current Biology, 34(19):4382–4396.e5.

Franke, K., Willeke, K. F., Ponder, K., Galdamez, M., Zhou, N., Muhammad, T., Patel, S., Froudarakis, E., Reimer, J., Sinz, F. H., and Tolias, A. S. (2022). State-dependent pupil dilation rapidly shifts visual feature selectivity. Nature, 610(7930):128–134. Number: 7930.

Gandhi, N. J. and Katnani, H. A. (2011). Motor functions of the superior colliculus. Annual Review of Neuroscience, 34:205–231.

Grosso, N. A. D., Graboski, J. J., Chen, W., Blanco-Hernández, E., and Sirota, A. (2017). Virtual Reality system for freely-moving rodents. Pages: 161232 Section: New Results.

Hayhoe, M. and Ballard, D. (2005). Eye movements in natural behavior. Trends in Cognitive Sciences, 9(4):188–194.

Holmgren, C. D., Stahr, P., Wallace, D. J., Voit, K.-M., Matheson, E. J., Sawinski, J., Bassetto, G., and Kerr, J. N. D. (2021). Visual pursuit behavior in mice maintains the pursued prey on the retinal region with least optic flow. eLife, 10:e70838.

Ito, S., Feldheim, D. A., and Litke, A. M. (2017). Segregation of Visual Response Properties in the Mouse Superior Colliculus and Their Modulation during Locomotion. The Journal of Neuroscience: The Official Journal of the Society for Neuroscience, 37(35):8428–8443.

Itokazu, T., Hasegawa, M., Kimura, R., Osaki, H., Albrecht, U.-R., Sohya, K., Chakrabarti, S., Itoh, H., Ito, T., Sato, T. K., and Sato, T. R. (2018). Streamlined sensory motor communication through cortical reciprocal connectivity in a visually guided eye movement task. Nature Communications, 9(1):338.

Krauzlis, R. J., Lovejoy, L. P., and Zénon, A. (2013). Superior colliculus and visual spatial attention. Annual Review of Neuroscience, 36:165–182.

Land, M. (2019). Eye movements in man and other animals. Vision Research, 162:1–7.

Land, M. F. (1999). Motion and vision: why animals move their eyes. Journal of Comparative Physiology. A, Sensory, Neural, and Behavioral Physiology, 185(4):341–352.

Land, M. F. and Tatler, B. W. (2009). Looking and Acting: Vision and Eye Movements in Natural Behaviour. Oxford University Press, Oxford.

Lima, A., Hou, Y., Beyeler, M., and Schneider, M. (2026). Beyond Neural Activity Prediction: Probing Latent Representations in Mouse V1 Digital Twins. arXiv:2605.23122 [q-bio.NC].

Mathis, A., Mamidanna, P., Cury, K. M., Abe, T., Murthy, V. N., Mathis, M. W., and Bethge, M. (2018). DeepLab-Cut: markerless pose estimation of user-defined body parts with deep learning. Nature Neuroscience, 21(9):1281–1289.

McCluskey, M. K. and Cullen, K. E. (2007). Eye, head, and body coordination during large gaze shifts in rhesus monkeys: movement kinematics and the influence of posture. Journal of Neurophysiology, 97(4):2976–2991.

Meyer, A. F., O’Keefe, J., and Poort, J. (2020). Two Distinct Types of Eye-Head Coupling in Freely Moving Mice. Current Biology, 30(11):2116–2130.e6.

Michaiel, A. M., Abe, E. T., and Niell, C. M. (2020). Dynamics of gaze control during prey capture in freely moving mice. eLife, 9:e57458.

Miura, S. K. and Scanziani, M. (2022). Distinguishing externally from saccade-induced motion in visual cortex. Nature, 610(7930):135–142.

Mongeau, J.-M. and Frye, M. A. (2017). Drosophila spatiotemporally integrates visual signals to control saccades. Current Biology, 27(19):2901–2914.e2.

Nishimoto, S., Vu, A. T., Naselaris, T., Benjamini, Y., Yu, B., and Gallant, J. L. (2011). Reconstructing visual experiences from brain activity evoked by natural movies. Current biology: CB, 21(19):1641–1646.

Parker, P. R., Abe, E. T., Beatie, N. T., Leonard, E. S., Martins, D. M., Sharp, S. L., Wyrick, D. G., Mazzucato, L., and Niell, C. M. (2022a). Distance estimation from monocular cues in an ethological visuomotor task. eLife, 11:e74708.

Parker, P. R. L., Abe, E. T. T., Leonard, E. S. P., Martins, D. M., and Niell, C. M. (2022b). Joint coding of visual input and eye/head position in V1 of freely moving mice. Neuron, 110(23):3897–3906.e5.

Parker, P. R. L., Martins, D. M., Leonard, E. S. P., Casey, N. M., Sharp, S. L., Abe, E. T. T., Smear, M. C., Yates, J. L., Mitchell, J. F., and Niell, C. M. (2023). A dynamic sequence of visual processing initiated by gaze shifts. Nature Neuroscience, 26(12):2192–2202. Number: 12 Publisher: Nature Publishing Group.

Salem, W., Cellini, B., Frye, M. A., and Mongeau, J.-M. (2020). Fly eyes are not still: A motion illusion in Drosophila flight supports parallel visual processing. Journal of Experimental Biology, 223(10):jeb212316.

Sato, T. R., Itokazu, T., Osaki, H., Ohtake, M., Yamamoto, T., Sohya, K., Maki, T., and Sato, T. K. (2019). Interhemispherically dynamic representation of an eye movement-related activity in mouse frontal cortex. eLife, 8:e50855.

Savier, E. L., Chen, H., and Cang, J. (2019). Effects of Loco-motion on Visual Responses in the Mouse Superior Colliculus. Journal of Neuroscience, 39(47):9360–9368.

Sharp, S. L., Shin, J., Martins, D. M., Jones, K., and Niell, C. M. (2025). Neural dynamics in superior colliculus of freely moving mice. Cell Reports, 44(10).

Skyberg, R. J. and Niell, C. M. (2024). Natural visual behavior and active sensing in the mouse. Current Opinion in Neurobiology, 86:102882.

Verdone, B. M., Chang, H. H. V., Roberts, D. C., and Cullen, K. E. (2026). Eye-head coordination during goal-directed orienting in mice. Communications Biology, 9(1):732.

Wallace, D. J., Greenberg, D. S., Sawinski, J., Rulla, S., Notaro, G., and Kerr, J. N. D. (2013). Rats maintain an overhead binocular field at the expense of constant fusion. Nature, 498(7452):65–69.

Wang, L., Liu, M., Segraves, M. A., and Cang, J. (2015). Visual experience is required for the development of eye movement maps in the mouse superior colliculus. Journal of Neuroscience, 35(35):12281–12286.

Xu, A., Hou, Y., Niell, C. M., and Beyeler, M. (2023). Multimodal Deep Learning Model Unveils Behavioral Dynamics of V1 Activity in Freely Moving Mice. In The Thirty-Seventh Annual Conference on Neural Information Processing Systems.

Yarbus, A. L. (1967). Eye Movements and Vision. Plenum Press, New York. Translated from Russian by Basil Haigh.

Zahler, S. H., Taylor, D. E., Wong, J. Y., Adams, J. M., and Feinberg, E. H. (2021). Superior colliculus drives stimulus-evoked directionally biased saccades and attempted head movements in head-fixed mice. eLife, 10:e73081.

Zahler, S. H., Taylor, D. E., Wright, B. S., Wong, J. Y., Shvareva, V. A., Park, Y. A., and Feinberg, E. H. (2023). Hindbrain modules differentially transform activity of single collicular neurons to coordinate movements. Cell, 186(14):3062–3078.e20.

